## Supporting information for "Pulmonary mesenchymal stem cells are engaged in distinct steps of host response to respiratory syncytial virus infection"

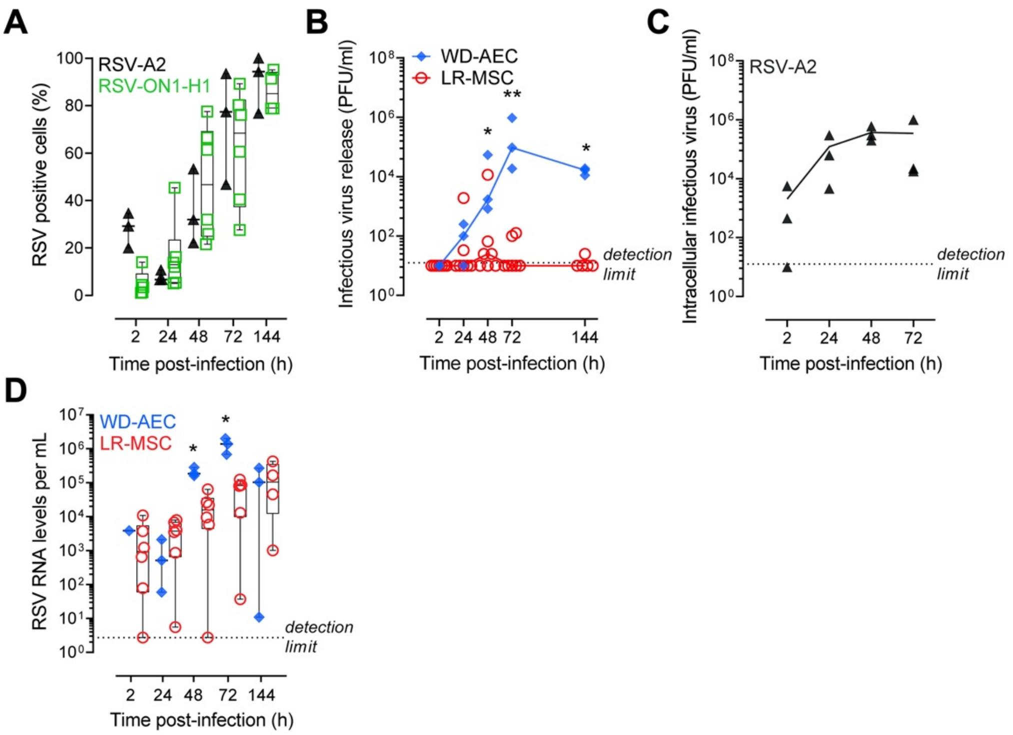


**Figure S1. Infection of pediatric LR-MSCs with RSV at a MOI of 1 PFU/cell**

(A) RSV F-protein positive LR-MSCs assessed by FCM and plotted over time. LR-MSCs were infected with 1 PFU/cell with RSV-A2 (n=3) or a clinical isolate RSV-ON1-H1 (n=4-6). Each symbol represents an individual donor. (B) Supernatants of infected LR-MSCs or apical washes of infected WD-AEC cultures were analyzed by a PFU assay. Cells were infected with RSV-ON1-H1 at a MOI of 1 PFU/cell. A Mann-Whitney U test was applied to compare the two cell types (WD-AECs, n=3 *versus* LR-MSCs, n=5-6). Each symbol represents an individual donor. *p<0.05, **p<0.01. (C) Intracellular infectious RSV titers in LR-MSCs infected with RSV-A2 at 1 PFU/cell. Each symbol represents an individual donor (n=3). (D) Extracellular viral RNA loads over time in supernatants of infected LR-MSCs or apical washes of infected WD-AEC cultures. Cells were infected with RSV-ON1-H1 at a MOI of 1 PFU/cell. A Mann-Whitney U test was applied to compare the two cell types (WD-AECs, n=3 *versus* LR-MSCs, n=4-6). Each symbol represents an individual donor. *p<0.05.

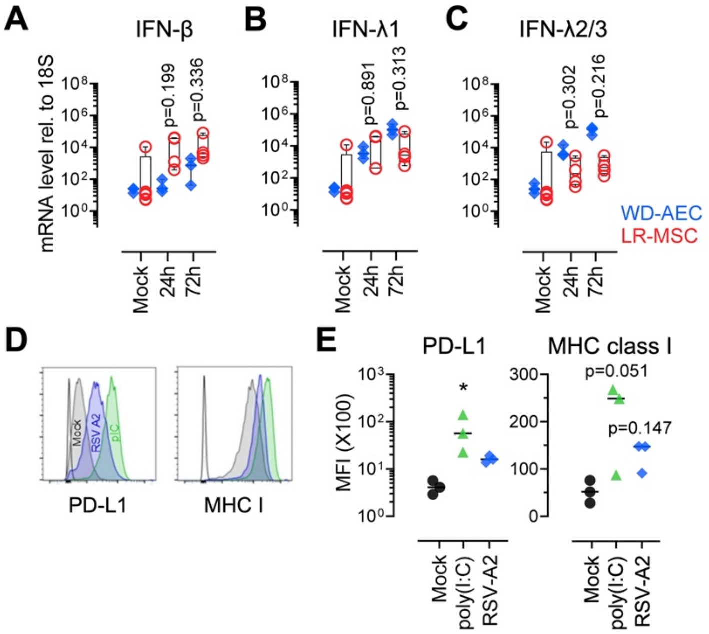


**Figure S2. IFN mRNA levels and PD-L1 and MHC class I surface expression in RSV-infected pediatric LR-MSCs**

(A-C) mRNA levels of IFN-β (A), IFN-λ1 (B), and IFN-λ2/3 (C) in LR-MSCs and WD-AECs infected with mock control or RSV-A2 at a MOI of 1 PFU/cell for 24 and 72 hours. Boxplots indicate the median value (centerline) and interquartile ranges (box edges), with whiskers extending to the lowest and the highest values. Each symbol represents an individual donor (LR-MSCs, n=4-6; WD-AECs, n=3). The data were compared with the Kruskal–Wallis test followed by Dunn's post hoc test. (D) Representative histogram of the surface expression of PD-L1 and MHC class I in pediatric LR-MSCs 24 h post-treatment with mock, poly(I:C) 10 μg/ml, and RSV-A2 at 1 PFU/cell. (E) Median fluorescence intensity (MFI) of PD-L1 and MHC class I expression. Each symbol represents an individual donor (n=3). The data were compared with the Kruskal–Wallis test followed by Dunn's post hoc test. *$p<0.05$.

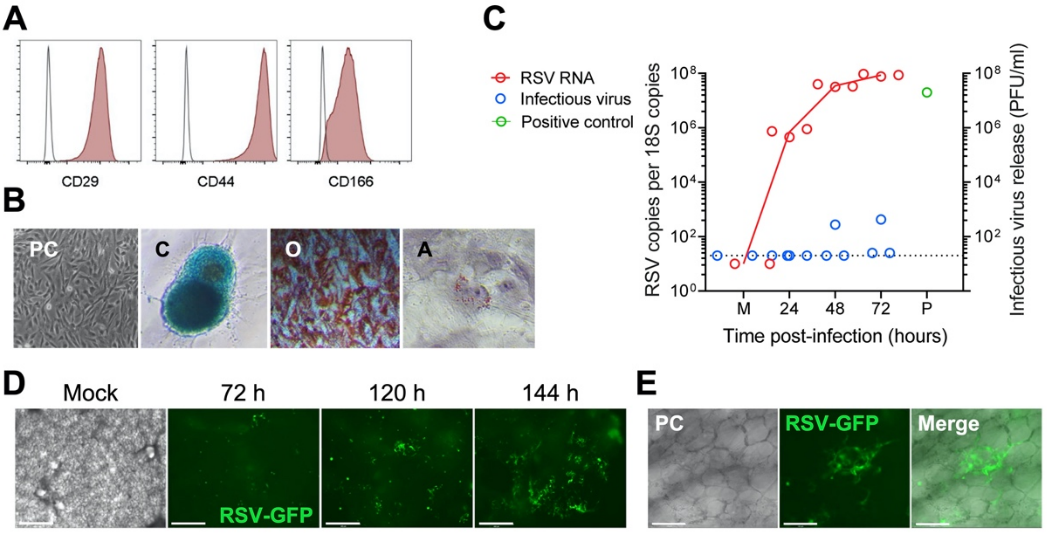


**Figure S3. RSV-infection of ovine PCLS and ovine LR-MSCs**

(A) Representative histograms showing expression of the surface markers CD29, CD44, and CD166. (B) Representative phase-contrast (PC) micrograph showing morphology in culture and demonstrates plastic adherence. Representative images of Toluidine blue, Alizarin Red S, and Oil Red O stainings after chondrogenic (C), osteogenic (O), and adipogenic (A) differentiation, respectively. Magnification 100X (PC, O), 200X (C, A). (C) Cell-associated RSV RNA loads expressed as RSV copies per $10^9$ 18S copies (red empty circles) and infectious virus release in PFU per ml (blue empty circles) over time following infection of primary ovine LR-MSCs with 0.1 PFU/cell of RSV-A2 determined 24, 48, and 72 h p.i. Each symbol represents an individual donor (n=3). The positive control (P) is the virus preparation used for the infections having a titer of $2x10^7$ PFU/ml. (M, mock). (D-E) Ovine PCLS have been infected with RSV-GFP at $5x10^5$ PFU per PCLS. Representative fluorescence micrographs are shown at 72, 120, and 144 hours p.i. Scale bar, 125 µm (D). Representative fluorescence micrographs indicating infection of pneumocytes. Scale bar, 650 µm (E).

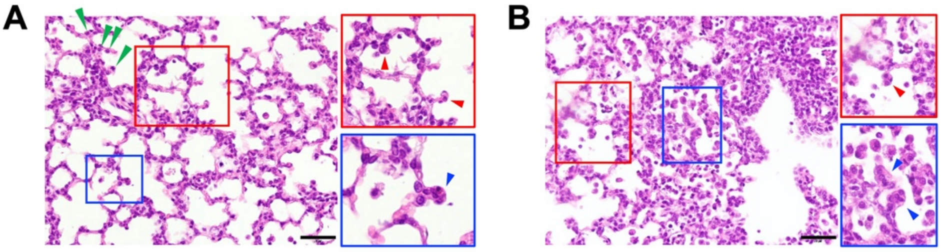


**Figure S4. Presence of potential syncytium in neonatal lungs following RSV infection**

(A, B) Representative histopathological sections of the lung tissue from lambs 6 days p.i. infected with RSV-A2. The colored boxes (red, blue) and arrowheads (red, blue) indicate the presence of potential syncytia and the green arrowheads indicate dome-shaped type 2 alveolar cells lining the alveolar wall, indicative for type 2 alveolar cell hyperplasia. Scale bar, 50 µm.

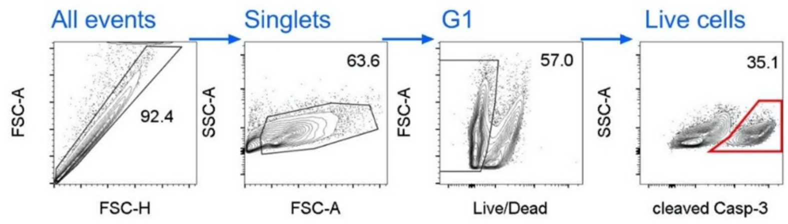


**Figure S5. FCM assay for apoptosis detection *in vivo***

Gating strategy for the detection of cleaved caspase-3-positive cells. (G1, gate 1).

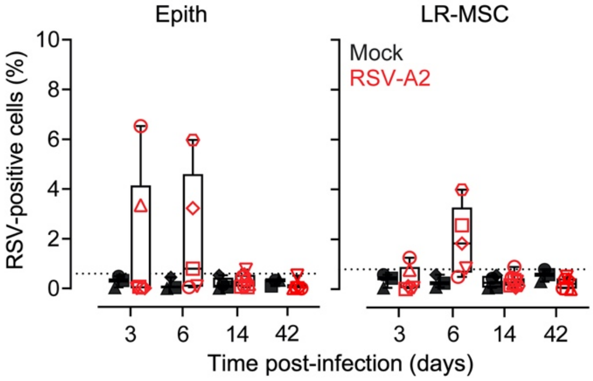


**Figure S6. Targeting of Epithelial cells and LR-MSCs following neonatal RSV infection**

RSV-positive epithelial cells (Epith) and LR-MSCs in lung cell suspensions were detected with an FCM assay 3, 6, 14, and 42 days following RSV-A2 infection of neonates. The dashed line depicts the detection limits (0.6% for Epith and 0.8% for LR-MSCs). Boxplots indicate the median value (centerline) and interquartile ranges (box edges), with whiskers extending to the lowest and the highest values. Each symbol represents an individual animal (mock, n=3-4; RSV, n=5-8).

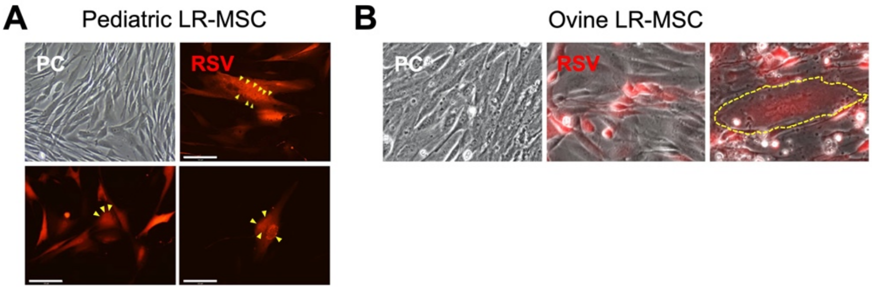


**Figure S7. Formation of syncytium upon RSV infection of human and ovine LR-MSCs**

(A, B) Representative micrographs of pediatric (A) and ovine (B) LR-MSCs infected with 0.1 PFU/cell of RSV-mCherry 48-72 hours p.i. Giant multinucleated cells, indicative of syncytium formation, were observed. Yellow arrowheads indicate clusters of nuclei and the yellow dashed line indicates a giant multinucleated cell. PC, phase-contrast. Magnification 100X (PC, pediatric LR-MSCs) and 400X (PC and RSV, ovine LR-MSCs). Scale bar, 125 µm.

**Table S1.** List of significant DEGs in RSV-infected *versus* mock-treated animals.

**RSV *versus* Ctrl 6 days p.i.**

| Gene ID | Gene Symbol | Log2 Fold Change | P value | $P_{adj}$ | Alternative name & function | Biological pathway | Ref. |
|---|---|---|---|---|---|---|---|
| ENSOARG00000020741 | TNFSF10 | 1.46 | 1.43E-09 | 1.10E-05 | TRAIL<br>pro-apoptotic | Apoptosis | (1, 2) |
| ENSOARG00000010042 | STC1 | 1.21 | 3.66E-06 | 1.00E-02 | Stanniocalcin-1<br>pro-apoptotic | Apoptosis | (3) |
| ENSOARG00000014648 | RSAD2 | 1.55 | 3.89E-11 | 7.44E-07 | Viperin<br>ISG | IFN pathway | (4) |
| ENSOARG00000007233 | ISG15 | 1.61 | 1.73E-09 | 1.10E-05 | ISG | IFN pathway | (5) |
| ENSOARG00000025182 | BST-2A | 1.40 | 3.38E-09 | 1.62E-05 | ISG<br>ISG | IFN pathway | (6) |
| ENSOARG00000013440 | IFI44 | 1.37 | 4.11E-08 | 1.57E-04 | ISG | IFN pathway | (7) |
| ENSOARG00000002881 | OASL* | 1.33 | 5.81E-07 | 1.85E-03 | ISG | IFN pathway | (8) |
| ENSOARG00000014800 | IFIT3 | 1.29 | 4.68E-06 | 1.12E-02 | RIG-G<br>ISG | IFN pathway | (9) |
| ENSOARG00000015177 | IFIT1 | 1.26 | 6.92E-06 | 1.32E-02 | ISG | IFN pathway | (10) |
| ENSOARG00000016787 | BST-2B | 1.03 | 6.53E-06 | 1.32E-02 | ISG | IFN pathway | (6) |
| ENSOARG00000001138 | HERC6* | 1.25 | 1.35E-05 | 2.15E-02 | Linked to ISG15 activation | IFN pathway | (11) |
| ENSOARG00000001815 | IRF* | 1.24 | 1.30E-05 | 2.15E-02 | IFN pathway | IFN pathway | (12) |
| ENSOARG00000013421 | IFI44L | 1.12 | 1.63E-05 | 2.40E-02 | ISG | IFN pathway | (7, 13) |
| ENSOARG00000014413 | IFI27L2* | 1.10 | 3.18E-05 | 4.35E-02 | ISG12b<br>ISG | IFN pathway | (14) |

ISG, interferon (IFN)-stimulated gene.

**RSV *versus* Ctrl 42 days p.i.**

| Gene ID | Gene Symbol | Log2 Fold Change | P value | $P_{adj}$ | Alternative name & function | Biological pathway | Ref. |
|---|---|---|---|---|---|---|---|
| ENSOARG00000009964 | TRPC4 | 1.15 | 4.84E-05 | 2.74E-02 | Calcium channel<br>Endothelial permeability<br>Vasodilation | Angiogenesis<br>Endothelium<br>Calcium | (15) |
| ENSOARG00000001060 | KDR | 1.10 | 1.54E-04 | 4.82E-02 | VEGF Receptor 2 (VEGFR2) | Angiogenesis<br>Endothelium<br>Calcium | (16) |
| ENSOARG00000003922 | MXRA8 | 0.99 | 7.96E-06 | 9.27E-03 | DICAM<br>Angiogenesis<br>Cartilage formation | Angiogenesis<br>Endothelium<br>Calcium | (17) |
| ENSOARG00000000498 | ITGA9 | 0.93 | 1.21E-04 | 4.49E-02 | Integrin $\alpha$9<br>Receptor of VCAM1 | Angiogenesis<br>Endothelium<br>Calcium | (18) |
| ENSOARG00000012316 | PTTG1IP* | 0.76 | 1.59E-04 | 4.85E-02 | Activator of FGF | Angiogenesis<br>endothelium<br>Calcium | (19) |
| ENSOARG00000012715 | PTGIS | 0.73 | 1.25E-04 | 4.49E-02 | Prostaglandin I2<br>Potent vasodilator<br>Inhibition of blood clot formation | Angiogenesis<br>Endothelium<br>Calcium | (20) |
| ENSOARG00000013623 | PPP3CA | -0.62 | 1.33E-04 | 4.49E-02 | Calcineurin<br>Calcium signaling<br>VEGF pathway | Angiogenesis<br>Endothelium<br>Calcium | (21) |

| | | | | | | | |
|---|---|---|---|---|---|---|---|
| ENSOARG00000006037 | BMPER | -0.91 | 7.99E-05 | 3.79E-02 | Inhibition of BMP function<br>Pro-angiogenic | Angiogenesis<br>Endothelium<br>Calcium | (22) |
| ENSOARG00000011962 | TRPM2 | -1.23 | 1.41E-05 | 1.50E-02 | Endothelial barrier function<br>Increase vascular permeability | Angiogenesis<br>Endothelium<br>Calcium | (23) |
| ENSOARG00000009518 | SOX4* | 0.64 | 7.50E-05 | 3.70E-02 | Associated with Wnt signaling | Differentiation<br>Development | (24) |
| ENSOARG00000011866 | PGAP6 | 0.62 | 1.33E-04 | 4.49E-02 | TMEM8A<br>Embryonic development | Differentiation<br>Development | (25) |
| ENSOARG00000009005 | C2CD3 | -0.54 | 8.82E-05 | 4.00E-02 | Associated with SHH signaling | Differentiation<br>Development | (26) |
| ENSOARG00000003978 | NOC3L | -0.55 | 7.41E-06 | 9.27E-03 | Adipogenesis | Differentiation<br>Development | (27) |
| ENSOARG00000019294 | ABCA12 | -1.13 | 3.81E-05 | 2.56E-02 | Lung development<br>ABCA12 KO leads to alveolar collapse | Differentiation<br>Development | (28) |
| ENSOARG00000020209 | CHPF | 0.86 | 5.82E-05 | 2.98E-02 | Chondroitin sulfate synthase-2<br>Regulation of ECM | ECM<br>Remodeling | (29) |
| ENSOARG00000008205 | IGSF8 | 0.66 | 1.16E-04 | 4.49E-02 | Negative regulator of TGF-beta | ECM<br>Remodeling | (30) |
| ENSOARG00000005315 | MMP1 | -1.06 | 1.43E-04 | 4.64E-02 | Breakdown of ECM | ECM<br>Remodeling | (31, 32) |
| ENSOARG00000005084 | MMP3 | -1.41 | 1.19E-06 | 2.17E-03 | Breakdown of ECM | ECM<br>Remodeling | (31, 32) |
| ENSOARG00000024340 | Metazoa_SRP | 1.85 | 2.47E-11 | 3.16E-07 | 7SL RNA<br>Non-coding RNA | Transcription regulation | (33) |
| ENSOARG00000023771 | miRNA* | 1.58 | 9.81E-09 | 4.19E-05 | Non-coding RNA | Transcription regulation | |
| ENSOARG00000022840 | RNase_MRP | 1.58 | 6.06E-10 | 3.88E-06 | Ribonuclease | Transcription regulation | (34) |
| ENSOARG00000022831 | 5_8S_rRNA | 1.53 | 7.01E-08 | 2.24E-04 | 5.8S ribosomal RNA<br>Non-coding RNA | Transcription regulation | |
| ENSOARG00000021831 | miRNA* | 1.43 | 3.88E-07 | 8.29E-04 | Non-coding RNA | Transcription regulation | |
| ENSOARG00000023915 | RNaseP_nuc | 1.34 | 2.66E-06 | 4.26E-03 | Ribonuclease | Transcription regulation | |
| ENSOARG00000023715 | 7SK RNA | 1.18 | 2.09E-05 | 1.81E-02 | Non-coding RNA | Transcription regulation | (35) |
| ENSOARG00000012889 | CPSF2 | -0.47 | 5.13E-05 | 2.74E-02 | mRNA processing and polyadenylation | Transcription regulation | (36) |
| ENSOARG00000017276 | MAGE2* | 1.18 | 1.76E-05 | 1.73E-02 | MAGE-like protein<br>Protein trafficking and recycling | Secretory pathway & cytoskeleton | (37) |
| ENSOARG00000005241 | MARCHF9 | 0.92 | 4.86E-05 | 2.74E-02 | Protein processing | Secretory pathway & cytoskeleton | (38) |
| ENSOARG00000018257 | TRIM3 | 0.59 | 3.92E-05 | 2.56E-02 | BERP<br>Interact with myosins | Secretory pathway & cytoskeleton | (39, 40) |
| ENSOARG00000019076 | TBC1D20 | 0.41 | 1.67E-04 | 4.97E-02 | Inhibition of RAB1<br>Autophagosome maturation | Secretory pathway & cytoskeleton | (41) |

| | | | | | | | |
|---|---|---|---|---|---|---|---|
| ENSOARG00000011359 | MAP7D3* | -0.35 | 1.14E-04 | 4.49E-02 | Regulates microtubule assembly and stability | Secretory pathway & cytoskeleton | (42) |
| ENSOARG00000011149 | CRYBG1 | -0.89 | 1.27E-04 | 4.49E-02 | AIM1<br>Suppresor of cell migration | Secretory pathway & cytoskeleton | (43) |
| ENSOARG00000006047 | PLS1 | -1.18 | 5.12E-05 | 2.74E-02 | Plastin-1/Fimbrin<br>Cross-linking with actin<br>Formation of filopodia | Secretory pathway & cytoskeleton | (44) |
| ENSOARG00000012318 | HMG20B | 0.77 | 2.12E-05 | 1.81E-02 | DNA repair<br>Mitosis | Other | (45) |
| ENSOARG00000013664 | SKA3 | -1.13 | 1.14E-04 | 4.49E-02 | Mitosis | Other | (46) |
| ENSOARG00000019275 | BARD1 | -1.22 | 2.85E-05 | 2.14E-02 | DNA repair | Other | (47) |
| ENSOARG00000009025 | ZNF503* | 0.98 | 6.22E-06 | 8.85E-03 | Fanconi anemia | Other | |
| ENSOARG00000009492 | COQ10A | 0.63 | 1.45E-04 | 4.64E-02 | Metabolism | Other | |
| ENSOARG00000020345 | WDR53 | -0.35 | 1.21E-04 | 4.49E-02 | Unknown function | Other | |
| ENSOARG00000000778 | MET | -0.78 | 9.06E-05 | 4.00E-02 | Receptor tyrosine kinase | Other | |
| ENSOARG00000001759 | FANCL | -0.89 | 4.00E-05 | 2.56E-02 | Fanconi anemia | Other | |
| ENSOARG00000004915 | TM4SF18 | -1.22 | 2.70E-05 | 2.14E-02 | Unknown function | Other | |
| ENSOARG00000019309 | ATIC | -1.34 | 2.95E-07 | 7.56E-04 | Inosine monophosphate synthase | Other | |

* Genes annotated manually.

**Table S2.** Patient characteristics

| Patient ID | Age (months) | Pathology |
|---|---|---|
| PL002 | 143 | CPAM |
| PL003 | 10 | Congenital lobar over inflation |
| PL004 | 153 | Chronic bronchiolitis/pneumonia |
| PL005 | 5 days | Bronchial atresia |
| PL006 | 11 | CPAM |
| PL021 | 6 | CPAM |
| PL012 | 1.5 | Lobar emphysema |
| PL018 | 181 | Aspergilloma |
| PL009 | 5 | CPAM |

CPAM, congenital pulmonary airway malformation.

**Table S3.** List of qPCR primers

| Gene | Primer | Sequence (5'-3') | Ref. |
|---|---|---|---|
| 18S rRNA | FW | CGCCGCTAGAGGTGAAATTC | |
| | RV | GGCAAATGCTTTCGCTCTG | (48) |
| | P | FAM-TGGACCGGCGCAAGACGGA-TAMRA | |
| IFN-β | FW | CGCCGCATTGACCATCTA | |
| | RV | TTAGCCAGGAGGTTCTCAACAATAGTGTCA | (49) |
| | P | FAM-TCAGACAAGATTCATCTAGCACTGGCTGGA-BHQ1 | |
| IFN λ1 | FW | GGACGCCTTGGAAGAGTCACT | |
| | RV | AGAAGCCTCAGGTCCCAATTC | (50) |
| | P | FAM-AGTTGCAGCTCTCCTGTCTTCCCCG-BHQ1 | |
| RIG-I | FW | CCAAGCCAAAGCAGTTTTCAA | |
| | RV | CACATGGATTCCCCAGTCATG | (51) |
| | P | FAM-TTGAAAAAAGAGCAAAGATATTCTGTGCCCGAC-TAMRA | |
| MDA5 | FW | GATTCAGGCACCATGGGAAGT | |
| | RV | AGGCCTGAGCTGGAGTTCTG | (51) |
| | P | FAM-GGGATGCTCTTGCTGCCACATTCTCTT-TAMRA | |
| 2',5'-OAS | FW | GATTCAGGCACCATGGGAAGT | |
| | RV | AGGCCTGAGCTGGAGTTCTG | (52) |
| | P | FAM-GGGATGCTCTTGCTGCCACATTCTCTT-TAMRA | |
| MxA | FW | CAGCACCTGATGGCCTATCAC | |
| | RV | CATGAACTGGATGATCAAAGG | (52) |
| | P | FAM-AGGCCAGCAAGCGCATCTCCAG-TAMRA | |
| Viperin | FW | CACAAAGAAGTGTCCTGCTTGGT | |
| | RV | AAGCGCATATATTTCATCCAGAATAAG | (52) |
| | P | FAM-CCTGAATCTAACCAGAAGATGAAAGACTCC-TAMRA | |
| ICAM-1 | FW | TGCAGACAGTGACCATCTACAGC | |
| | RV | TCTGAGACCTCTGGCTTCGTC | (53) |
| | P | FAM-TTCCGGCGCCCAACGTGATT-TAMRA | |
| Nucleolin | FW | TCGCGAAGGCAGGTAAAAA | |
| | RV | CGACCTCTTCTCCACTGCTATCA | (54) |
| | P | FAM-AAGGTGACCCCAAGAAAATGGCTCCTC-TAMRA | |
| Annexin A2 | FW | GTGAAGAGGAAAGGAACCGA | |
| | RV | CTTGATGCTCTCCAGCATGT | (55) |
| | P | - | |
| TLR4 | FW | CAGAGTTGCTTTCAATGGCATC | |
| | RV | AGACTGTAATCAAGAACCTGGAGG | (56) |
| | P | - | |
| CX3CR1 | FW | AGTGTCACCGACATTTACCTCC | |
| | RV | AAGG CGGTAGTGAATTTGCAC | (57) |
| | P | - | |
| hRSV-A2 (L) | FW | GAACTCAGTGTAGGTAGAATGTTTGCA | |
| | RV | TTCAGCTATCATTTTCTCTGCCAAT | (58) |
| | P | FAM-TTTGAACCTGTCTGAACATTCCCGGTT-TAMRA | |
| RSV-A (N) | FW | TGCTAAGACYCCCCACCGTAAC | |
| | RV | GGATTTTTGCAGGATTGTTTATGA | (59) |
| | P | C5CT6GC7CT87W7CA-BHQ1 | |

FW, forward; RV, reverse; P, probe.

**Table S4.** List of antibodies used

| Antibody | Dilution | Host, isotype | Clone | Reference, Source |
|---|---|---|---|---|
| Anti-CD73 | 1:200 | Mouse, IgG1 | AD2 | ab30451, abcam |
| Anti-CD90 | 1:100 | Mouse IgG1 | 5.00E+10 | 555593, BD Pharming |
| Anti-CD105, FITC | 1:25 | Mouse, IgG2a | MEM-229 | ab53318, abcam |
| anti-RSV F protein | 1:100 | Mouse, IgG2a | 131-2A | MAB8599, Millipore |
| Anti-RSV, biotin | 1:160 | Goat, IgG | - | 7950-0104, Bio-Rad |
| Cleaved Caspase-3 | 1:800 | Rabbit | Asp175 | 9661, Cell Signling |
| Anti-CD29 | 1:100 | Mouse, IgG1 | S-FW4-101 | S-BOV2034 WSU |
| Anti-CD44 | 1:100 | Mouse, IgG3 | S-BAG40A | S-BOV2037WSU |
| Anti-CD166, FITC | 1:20 | Mouse, IgG1 | 3A6 | MCA1926FT, Bio-Rad |
| Anti-CD31, FITC | 1:40 | Mouse, IgG2a | CO.3E1D4 | MA1-80360,Thermofisher |
| Anti-CD45, PE | 1:100 | Mouse, IgG1 | 1.11.32 | MCA2220PE, Bio-Rad |
| Anti-pan-cytokeratin, APC | 1:800 | Mouse, IgG1 | C-11 | MA1-10325, Thermofisher |
| Anti-Mouse IgG1, af488 | 1:2000 | Goat, IgG | - | A-21121, Thermofisher |
| Anti-Mouse IgG1, PE | 1:200 | Goat, IgG | - | A-21129, Thermofisher |
| Anti-Mouse IgG1, PerCP-Cy5.5 | 1:80 | Rat, IgG | RMG1-1 | 406612, Biolegend |
| Anti-Mouse IgG1, PE-Cy7 | 1:640 | Rat, IgG | RMG1-1 | 406614, Biolegend |
| Anti-Mouse IgG1, af647 | 1:1000 | Goat, IgG | - | A-21240, Thermofisher |
| Anti-Mouse IgG1, BV421 | 1:80 | Rat, IgG | RMG1-1 | 406616, Biolegend |
| Anti-Mouse IgG3, PerCP/Cy5.5 | 1:200 | Goat, IgG | - | 1100-13, Southernbiotech |
| Streptavidin, Pacific Blue | 1:200 | - | - | S11222, Thermofisher |
| Anti-Mouse IgG2a, af647 | 1:1000 | Goat, IgG | - | A-21241, Thermofisher |
